# A transcriptome atlas of pea seed development guides the identification of PsLEC1-like as a key regulator of seed size

**DOI:** 10.64898/2026.05.15.725475

**Authors:** Yara Noureddine, Titouan Bonnot, Christine Le Signor, Johanne Thevenin, Jérôme Verdier, Nadia Rossin, Myriam Sanchez, Jonathan Kreplak, Marion Dalmais, Karine Gallardo-Guerrero, Bertrand Dubreucq, Vanessa Vernoud

## Abstract

Grain legumes such as pea (*Pisum sativum* L.) accumulate large amounts of seed storage proteins without nitrogen fertilization due to their symbiosis with nitrogen-fixing bacteria, making them a key source of plant-based proteins. Seed growth and the accumulation of seed storage proteins are tightly regulated by complex gene networks; however, the mechanisms governing these processes in pea remain poorly understood. In this study, we generated a comprehensive seed expression atlas covering six developmental stages in pea (cv Caméor), including the key transition stage from embryogenesis to early seed filling, providing a detailed temporal resolution of transcriptional dynamics throughout seed development in this species. Co-expression network analysis highlighted several candidate transcription factors potentially involved in the transition towards seed filling. Among them, we characterized the seed-specific NF-YB transcription factor PsLEC1-like (PsL1L), the major LEC1-type factor expressed during early pea seed development. Functional analyses using TILLING mutants demonstrated that loss of *PsL1L* function reduces seed size and seed nitrogen content and impairs early embryo growth from the end of embryogenesis. Finally, we show that the expression of the B3-domain transcription factor *PsFUS3*, but not that of *PsLEC2* or *PsABI3*, is reduced in the loss-of-function *l1l* mutant, suggesting that PsL1L acts upstream of PsFUS3 to control seed size.

## Introduction

Grain legumes such as pea (*Pisum sativum* L.) can help meet the growing demand for plant proteins in food and feed. They accumulate large quantities of proteins in their seeds (23% in pea, (Burstin et al., 2011) even in the absence of nitrogen (N) fertilizer, due to the unique ability of their root system to form symbioses with soil bacteria (genus *Sinorhizobium*) that are able to fix atmospheric nitrogen in dedicated structures called nodules (Lindström & Mousavi, 2020). The use of legumes in cropping systems thus reduces greenhouse gas emissions from fertilizer production and use, and increases soil fertility (Nicastro & Poór, 2022) This makes legumes an attractive source of plant protein, providing nutritional benefits while supporting lower environmental impacts.

Pea seed development is classically divided into three successive phases: embryogenesis, seed filling corresponding to the accumulation of reserves (proteins and starch) in the cotyledons, and maturation (Weeden, 1995). Embryogenesis is characterized by intense mitotic activity, during which the surrounding tissues, the endosperm and seed coat, play a key role in nutrient supply to the growing embryo. Importantly, in legumes the final number of cotyledon cells is established at the end of embryogenesis, and correlated with the final individual seed weight/size at maturity (Lemontey et al., 2000; Munier-Jolain & Ney, 1998). The transition to the filling phase occurs around 10-12 days after pollination (DAP), when seed water content drops below approximately 85%, and marks a shift from cell division-driven growth to growth predominantly driven by cell expansion (Schiltz et al., 2004). The filling phase is then characterized by the active accumulation of storage reserves, mainly proteins and starch, leading to a rapid increase in seed biomass. Finally, around 30-35 DAP, seeds undergo a period of intense desiccation associated with a reduction in metabolic activity, leading to a state of metabolic quiescence.

The seed storage proteins (SSP) in pea seeds determine their nutritional value. The majority are globulins, accounting for about 70% of total seed protein (Boulter and Croy, 1997). Other minor proteins accumulating in pea seeds include 2S albumins, which are rich in sulfur-containing amino acids (sulfur-AA). Globulins are encoded by multigene families and have been classified into 7S (vicilins and convicilins) and 11S (legumes) types according to their sedimentation coefficients after ultracentrifugation (Osborne, 1924). Convicilins, which account for only 6% of total seed protein, contain higher levels of sulfur-AA than vicilins (Bourgeois et al., 2009; Tzitzikas et al., 2006). Seed storage protein accumulation in legumes is primarily regulated at the transcriptional level (Verdier & Thompson, 2008). In pea the rate of accumulation of 7S and 11S globulins is strongly correlated with the expression levels of the genes encoding these proteins (Evans et al., 1984; Higgins et al., 1986). Consequently, the total amount of storage proteins largely depends on transcript abundance. In addition, post-translational modifications and processes occurring during protein transport can further influence the final abundance and composition of storage proteins. The various aspects of seed development, from embryogenesis to maturation are largely regulated by numerous transcription factors (TFs) (Verma et al., 2022).

In *Arabidopsis thaliana*, seed reserve accumulation is transcriptionally regulated by a set of master regulators collectively known as the LAFL network, comprising the B3-domain transcription factors LEAFY COTYLEDON 2 (AtLEC2), ABSCISIC ACID INSENSITIVE 3 (AtABI3), and FUSCA3 (AtFUS3), as well as the NF-YB factor LEAFY COTYLEDON 1 (AtLEC1) (reviewed in Lepiniec et al., 2018). These regulators control the expression of genes involved in seed maturation, including storage proteins and lipid biosynthesis genes, acting alone or in combination with other transcription factors such as bZIPs (Baud et al., 2016; Fatihi et al., 2016). They form a complex regulatory network that coordinates the transition from embryogenesis to seed maturation and later to germination (Boulard et al., 2017). Likewise, genes in the LAFL family are subjected to strong transcriptional and post-transcriptional regulation (Lepiniec et al., 2018). Analyses of interactions among these master regulators suggest that *AtLEC1* acts at the highest level in the regulatory hierarchy controlling the maturation phase (Jo et al., 2019; Kagaya et al., 2005). Loss-of-function of *AtLEC1* causes defects in storage protein and lipid accumulation, acquisition of desiccation tolerance, and the suppression of germination and leaf primordia initiation (reviewed in Jo et al., 2019) and reduction in somatic embryogenesis. Among NF-YB family members, AtLEC1-LIKE (*AtL1L*) is the closest homolog of *AtLEC1.* It can partially complement the *lec1* mutations when ectopically expressed, and RNAi lines display embryo defects (Kwong et al., 2003) but, compared to *AtLEC1,* its function remains less clear.

In pea, the molecular mechanisms underlying seed development, including the control of seed size and seed filling, remain poorly understood. In this context, we recently generated proteomic data for early seed development (cv. Caméor) and analyzed the proteome reprograming of developing pea seeds (Henriet et al., 2021). However, a comprehensive transcriptomic landscape spanning the entire course of seed development is still lacking, despite the availability of its genome sequence (Kreplak et al., 2019), including an improved version of the reference cultivar Caméor (Kreplak et al., 2026).

In this study, we characterized the highly dynamic transcriptional landscape underlying pea seed development using RNA-seq analysis. Through co-expression network analysis and transcription factor (TF) mining, we identified key TFs potentially involved in the regulation of seed-related traits, including seed size and seed protein content and composition. Members of the NF-YB TF family were particularly prominent among those identified. Using a TILLING (Targeting Induced Local Lesions IN Genomes) mutant population (Dalmais et al., 2008), we performed functional analyses of *PsL1L* that revealed the key role of this gene in controlling embryo and seed growth.

## Materials and methods

### Plant Material and growth Conditions

Pea plants (*Pisum sativum* L., cv. Caméor), including the TILLING mutants analyzed in this study, were cultivated individually in 2-L pots containing a mixture of attapulgite and clay balls (1:3, v/v) and automatically irrigated with a nutrient solution containing 10:10:10 N:P:K. Growth conditions were maintained at a day/night temperature of 19/15 °C under a 16-h photoperiod with artificial lighting (250 µmol m⁻² s⁻¹).

### Seed sampling and seed tissue preparation

Emerging pea flowers were tagged on the day of pollination (closed flower stage 0.2, Maurer et al., 1965). Tagged pods were harvested at defined developmental stages, immediately placed on ice, and processed according to downstream analyses.

For transcriptomic analysis, pods from flowering nodes 2 or 3 were harvested at 8, 12, 16, 19, 23, and 29 days after pollination (DAP). For each pod, two seeds were used to determine mean individual seed fresh weight (FW), dry weight (DW; after drying at 80 °C for 48 h), and water content calculated as [(FW – DW) / FW] × 100. The remaining seeds were pooled, snap-frozen, and stored at −80 °C until RNA extraction. Four biological replicates were performed per developmental stage, each consisting of seeds from two pods.

For gene expression analysis in seed tissues (Figure 3D), Caméor seeds were collected at 10, 12, 14, and 16 DAP and dissected on ice. Liquid endosperm was collected using a P10 pipette, pooled, and immediately snap-frozen; it was no longer present at 16 DAP. Seed coats and embryos were rinsed twice in distilled water to remove residual endosperm, gently blotted dry, and snap-frozen. Three biological replicates were prepared per stage, each consisting of approximately 20 dissected seeds.

For the kinetic experiment (Figure 6A), pods from wild-type and *l1l-1* TILLING mutant plants were collected at 8, 12, 16, 20, 25, 30, 35, and 38 DAP. For each genotype and developmental stage, 6 to 8 pods containing 4 to 7 seeds each were analyzed. Individual seed FW, DW and water content were determined as described above.

### RNA extraction

Total RNA was extracted from ground seeds or seed tissues using the Spectrum™ Plant Total RNA kit (SIGMA). Trace of genomic DNA were removed using the One-Column DNase Digest Set (SIGMA) following the manufacturer’s instructions. RNA concentration and quality were evaluated using a NanoDrop™ 2000 spectrophotometer (Thermo Scientific) and an Agilent 2100 Bioanalyzer (Agilent Genomics) for samples intended for RNA sequencing.

### RNA sequencing

RNA-Seq libraries were prepared using an Illumina TruSeq Stranded mRNA sample prep kit, with 11 PCR cycles for library amplification. After library quality check on a Fragment Analyzer (Agilent Genomics), sequencing was performed on the Illumina HiSeq3000 to obtain 2×150 bp paired-end reads. After sequence quality assessment (FastQC v0.11.2 software, https://www.bioinformatics.babraham.ac.uk/projects/fastqc/), adapter and low-quality sequences were trimmed (Trimmomatic v0.32,Bolger et al., 2014). Paired reads were then mapped to the *P. sativum* v2 reference genome (Kreplak et al., 2026), and counted using nf-core rnaseq pipeline by default.

### Statistical analysis of transcriptomic data

Statistical analyses were performed with the R program v. 4.4.1 (R Core Team, 2022). Genes with at least 10 reads total in the experiment were filtered and considered for downstream analyses (25577 genes). Differential expression analysis was performed from raw counts with the DESeq2 package (Love et al., 2014). First, a Likelihood Ratio Test (LRT) was performed to determine significant differences in gene expression across seed development. Second, pairwise comparisons were used to compare time points. Genes with *P* < 0.05 after FDR correction using the Benjamini-Hochberg procedure (Benjamini & Hochberg, 1995), for both the LRT analysis and for at least one time point comparison were considered as significant.

For exploratory analyses described above, raw counts were normalized using the variance stabilizing transformation (VST) provided by ‘DESeq2’ v. 1.46.0 (Love et al., 2014). PCA was performed and visualized using ‘ade4’ (v. 1.7-24) and ‘factoextra’ (v. 2.0.0) R packages (Kassambara & Mundt, 2020; Thioulouse et al., 1997). Hierarchical clustering analysis was performed with ‘pheatmap’ v. 1.0.13 (Kolde, 2015), using the euclidean distance as dissimilarity measure (Supplementary Dataset S2). The number of clusters were determined manually after multiple testing. Gene Ontology (GO) enrichment analysis was performed by cluster with ClusterProfiler v. 4.16.6 (Yu et al., 2012), using genes identified as differentially expressed – and thus used for clustering – as reference. GO terms with *P* < 0.05 after FDR correction using the Benjamini-Hochberg procedure (Benjamini & Hochberg, 1995), and with a fold enrichment > 1 were considered as over-represented in the cluster compared to the reference. GO enrichment results are presented in supplementary Dataset S3.

### Analysis of transcription factors

To identify pea TFs, iTAK was run on proteins of the v2 pea genome annotation, using default parameters (Zheng et al., 2016). To identify developing seed-specific TFs, the gene expression atlas published in (Alves-Carvalho et al., 2015) was used. In this dataset, two tissues/conditions correspond to seed: seeds collected at 12 days after pollination (seed_12dap) and after five days of imbibition. First, TFs with TPM > 1 in at least one tissue/condition were filtered. Second, SPM scores were calculated for each gene, corresponding to TPM in seed_12dap/sum (TPM in all tissues/conditions). The higher the SPM value (between 0 and 1), the higher the specificity of gene expression in developing seed is. TFs in the top 5% of SPM values and with TPM > 2 in seed_12dap were considered as specifically expressed in developing pea seeds. Data are presented in supplementary Dataset S4.

### Co-expression network analysis

Weighted Gene Co-expression Network Analysis was conducted from RNA-Seq data (VST-transformed counts) of differentially expressed genes across seed development (20144 genes), using the R package ‘WGCNA’ v. 1.74 (Langfelder & Horvath, 2008). Adjacency matrix was built using a soft threshold power of 30. The minimum module size was set at 50 for module detection. Co-expression networks were then visualized with CYTOSCAPE v. 3.10.1 (Smoot et al., 2011), using a Prefuse Force Directed Layout, after filtering links with weight > 0.15. Data are presented supplementary Dataset S5.

### Phylogenetic analyses

Multiple sequence alignment was performed using ClustalW and the phylogenetic tree was constructed with MEGA12 (Kumar et al., 2024) using the Maximum Likelihood approach, with 1000 bootstrap.

### Screening for *PsL1L* TILLING mutants

The *l1l-1* and *l1l-2* mutants were identified by screening the Caméor TILLING population (Dalmais et al., 2008). PCR and Cel1 digestion were performed as previously described (Dalmais et al., 2008). A first amplicon of 925pb was amplified with primers L1L_N1F and L1L_N1R at an annealing temperature of 58 °C (Supplementary Table S1 for primer sequences). Nested PCR using this first amplicon as a template was carried out (870pb, annealing temperature 58 °C) with primers L1L_N2F and L1L-N2R labelled at the 5′ end with infra-red dyes IRD700 or IRD800, respectively (Biolegio). To confirm mutations, PCR amplification products were sequenced (Sanger sequencing, Genewiz) and analysed (Chromas Lite v.2.3software).

For each allele, M3 TILLING lines were back-crossed twice to Caméor in order to reduce the number of background mutations, and the plants phenotyped were progenies issued from at least two generations of selfing. All progenies were genotyped with specific dCAPS markers designed with the dCAPS Finder 2.0 program. The primer sequences and enzymes used for genotyping are available in Supplementary Table S1.

### Seed clearing for mutant phenotyping

Embryos were visualized using a BABB-based tissue-clearing method (Cabrera et al., 2018) optimized for pea seeds. Seeds from wild type and *l1l-1* mutant plants were collected at 8, 10 and 12 DAP using previously described procedures. Seeds were fixed in 3% glutaraldehyde in phosphate-buffered saline (PBS) under moderate vacuum for 1h, with pressure cycles applied every 15 min to enhance fixative penetration. Samples were then stored in fresh fixative at 4 °C for several days. After fixation seeds were rinsed twice in PBS and dehydrated through a graded ethanol series (30%, 50%, 70% and 90%, 30 min each), followed by incubation in absolute ethanol for 48 h at 4 °C. For clearing, samples first incubated in a 1:2 ethanol:BABB solution for 1 h and then transferred to 100% BABB until fully transparent. Seeds collected at 6-10 days after pollination (DAP) typically cleared within about three days, whereas older seeds (12-14 DAP) could require up to a week. Cleared seeds were mounted in BABB-filled imaging chambers and imaged using a Leica SP8 confocal microscope (excitation at 480 nm and emission at 500–600 nm). The natural autofluorescence produced by glutaraldehyde fixation provided strong contrast, allowing clear visualization of internal seed structures, including the embryo. Embryo perimeter was quantified by manually segmentation of embryo outlines in LAS X software (Version 5.1.0). To ensure imaging in the same plane across different seeds, samples were focused along the z-axis using a consistent anatomical reference point. Z-stacks were acquired with identical start and end planes, allowing selection of comparable optical sections between samples. Imaging parameters (laser power, pinhole, and detector gain) were kept constant.

### Quantitative RT-PCR

Two µg of purified RNA were reverse-transcribed with the iScript cDNA synthesis kit (Bio-Rad) according to manufacturer’s protocol in a final volume of 20 µl. Real-time PCR was performed on a LightCycler 480 (Roche) as described by Noguero et al., 2015 with GoTaq qPCR Master Mix (Promega). *PsEF1α (Psat.cameor.v2.1g212000)* and *PsHKG1 (Psat.cameor.v2.3g295150)* encoding a 5S ribosomal protein were used for the normalization of qRT-PCR data. Both genes were identified as stably expressed during seed development based on RNA-seq data generated in this study. For analyses specifically performed on seed tissues (Figure 3B), a third reference gene, *PsACTIN* (Psat.cameor.v2.5g339550), was also included. Relative expression levels were calculated using the ΔΔCt method described by Schmittgen and Livak (2008). Primer sequences for both target and reference genes are listed in Supplementary Table S1.

### Quantification of transcriptional activity in moss protoplasts

The interaction of transcription factors on a target promoter was analyzed by transient expression assays in *Physcomitrella patens* moss, as previously described by Thévenin et al., 2016. Transcription factors were cloned into the pBS TPp-A vector allowing their expression under the control of the rice ACTIN promoter. PsL1L wild-type and mutant sequences (L1L-1 and L1L-2) were synthesized with the flanking attB1 and attB2 recombination sites (Twist Bioscience), cloned into the pDONR207 entry vector using the Gateway technology, and subsequently recombined into the pBS TPp-A destination vector (Thévenin et al., 2012). A 250 bp fragment of the OLEOSIN 1 promoter was recombined into the pBS TPp-B vector, which drives GFP expression (Baud et al., 2016). Briefly, moss protoplasts were transfected with 5 µg of each plasmid, including one or more TF constructs (proACTIN::TF) and the reporter construct containing the target promoter fused to GFP (pOLE1::GFP). GFP fluorescence in moss protoplasts was quantified 48 h after transfection using a flow cytometry (Partec CyFlow Space, laser 488 nm, 20 mW) with SYSMEX (France) reagents. Fluorescence signals were collected within a defined gate (R4-Mean). The fluorescence was directly correlated to the transcriptional activity of the TF on the target promoter.

### Statistical analysis

For transcriptional activity in moss protoplasts statistical significance was assessed using a Kruskal–Wallis non-parametric test followed by Dunn’s post hoc test with Benjamini–Hochberg correction (α = 0.05), as described by Salaün (2025). Pairwise comparisons between wild type and *l1l* TILLING lines were performed using Student’s t-test or Mann–Whitney U test after testing for normality using the Shapiro–Wilk test. Analyses were performed in Rstudio (version 2026.1.0.392).

## Results

### Identification of genes differentially expressed across seed development

To characterize the transcriptome dynamics underlying pea seed development, we analyzed RNA-seq data at six developmental stages (8, 12, 16, 19, 23 and 29 DAP). Seed developmental stages were defined according to seed water content (Ney et al., 1993; Supplementary Table S2). The 8 DAP stage corresponded to the embryogenesis phase, characterized by low dry matter accumulation and high water content (>85%). At 12 DAP, seed water content dropped below 85%, marking the transition to the filling stage, which lasted up to 23 DAP. At 29 DAP, water content fell below 55%, corresponding to the beginning of the desiccation phase. A total of 25,577 genes were considered as expressed during seed development (Supplementary Dataset S1). Principal Component Analysis (PCA) revealed strong clustering of biological replicates, and clear separation between developmental stages, with PC1 separating embryogenesis (8 DAP) and late maturation (29 DAP) phases (Figure S1). Differential expression analysis identified 20,144 genes (79% of expressed genes) genes with significant temporal expression changes (see Methods). Interestingly, PC3 clearly separates 8 and 12 DAP, suggesting substantial changes at the transition between embryogenesis and seed filling (Figure S1).

Hierarchical clustering analysis of differentially expressed genes (DEGs) revealed six clusters with distinct temporal profiles corresponding to embryogenesis (cluster 4), transition between embryogenesis and seed filling (cluster 1), seed filling (clusters 6 and 5), late maturation (cluster 3), and embryogenesis/late maturation (cluster 2, Figure 1, A and B, supplementary Dataset S2). Gene Ontology (GO) enrichment analysis revealed that genes expressed during embryogenesis and at the transition with seed filling were associated with ‘DNA conformation change’ and ‘DNA replication’, ‘cytoskeleton organization’ and ‘cell wall biogenesis’ (Figure 1C, supplementary Dataset S3), consistent with cell proliferation and growth during early seed developmental stages. Genes expressed during seed filling are enriched for ‘photosynthesis’, ‘seedling development’, ‘carbohydrate homeostasis’, ‘cellular response to nitrogen levels’ and ‘regulation of lipid metabolic process’, highlighting coordination between energy production, metabolism and development. Late maturation phase is characterized by genes playing a role in RNA processing pathways, protein transport and modification (‘protein import’, ‘protein ubiquitination’, ‘protein dephosphorylation’), ‘negative regulation of cellular metabolic process’ and ‘regulation of response to water deprivation’, consistent with desiccation tolerance and the entry into quiescence (Figure 1C; supplementary Dataset S3).

**Figure 1.**
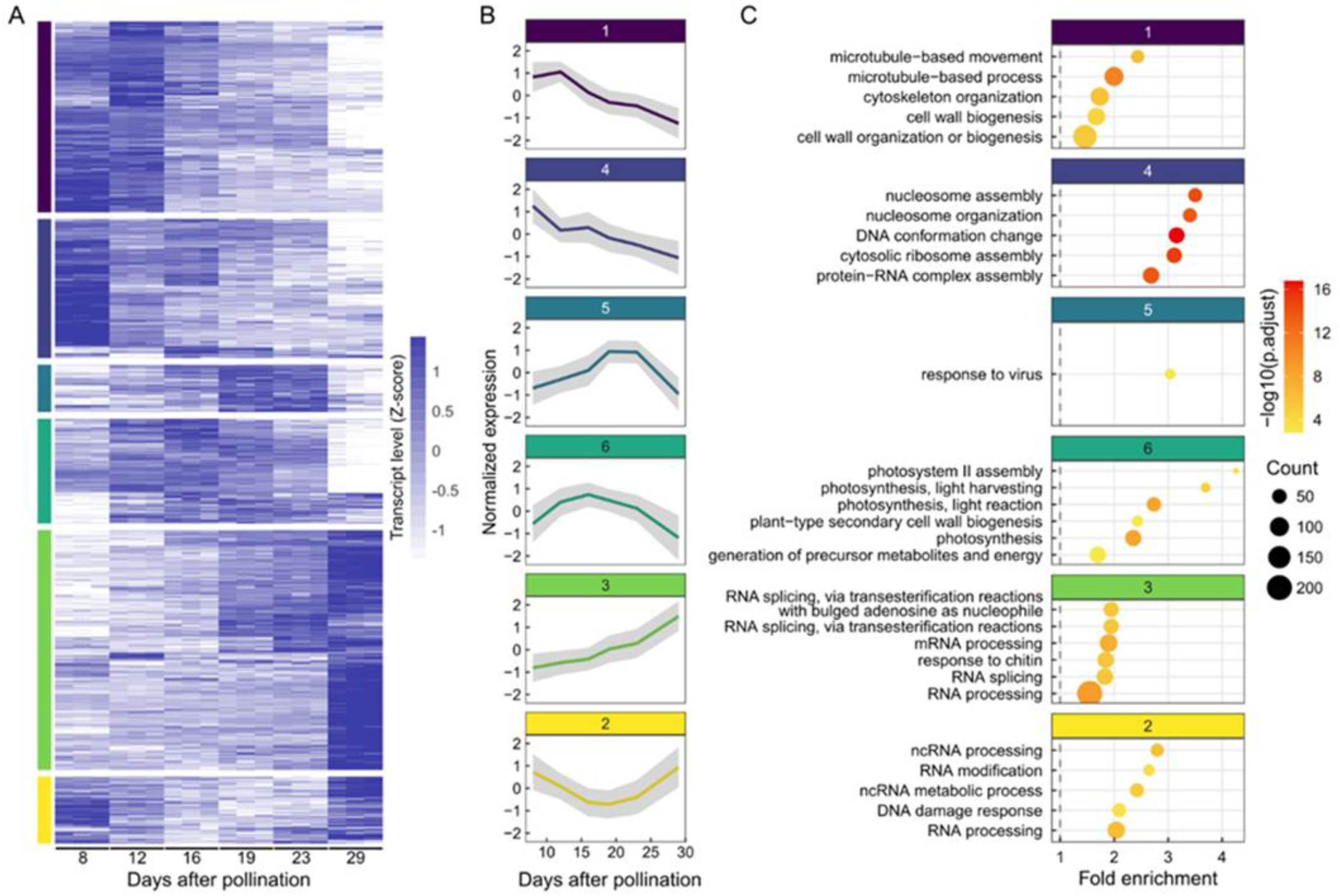
Clustering analysis of differentially expressed genes across pea seed development. (A) Heatmap representing clustering of differentially expressed genes (DEGs) across seed development. Data are scaled by row. (B) Average gene expression profiles of identified clusters. Lines represent mean values and shaded areas represent standard deviations. (C) Enriched biological processes in identified clusters. For each cluster, the top 5 enriched Gene Ontology (GO) terms (lowest adjusted P-values) with at least five genes were selected. Six GO terms are represented for clusters 6 and 3, due to equal adjusted P-values for two GO terms. The adjusted P-values were log10-transformed to help with visualization.

### Candidate transcription factors for regulating the transition towards seed filling

To further investigate temporal coordination of gene expression during pea seed development, we built a co-expression network from the set of DEGs. This network revealed distinct gene groups corresponding to the expression clusters identified above, and highlighted the continuum between genes expressed from early to late developmental stages (Figure 2A). To identify putative regulatory genes of pea seed filling, we focused on transcription factors (TFs) and seed storage protein (SSP) genes. The TFs subnetwork in which we included genes encoding SSP (TF-SSP subnetwork) evidenced two main groups of highly co-expressed TFs: those expressed at early stages (8 and 12 DAP) and those at late stages (29 DAP, Figure 2B). Next, we reasoned that seed-specific TFs expressed at the transition between embryogenesis and seed filling (12 DAP) are candidate regulators for the induction of SSP synthesis. We first identified TFs with specific expression in developing seeds, using a gene expression atlas for pea (Alves-Carvalho et al., 2015; see methods and supplementary Dataset S4). We then filtered our network in Figure 2B to build a network composed of 52 seed-specific TFs (Figure 2D). Most of these TFs showed peak expression at 8 DAP (30 TFs) or at 12 DAP (19 TFs), and include members of the B3, C2H2, HSF, MADS-M-type, MYB and NF-YB families (Figure 2, C and D). We found that three of the four NF-YB exhibited a peak of expression at 12 DAP (Figure 2, D and E), suggesting a role in preparing the seed for storage processes.

**Figure 2.**
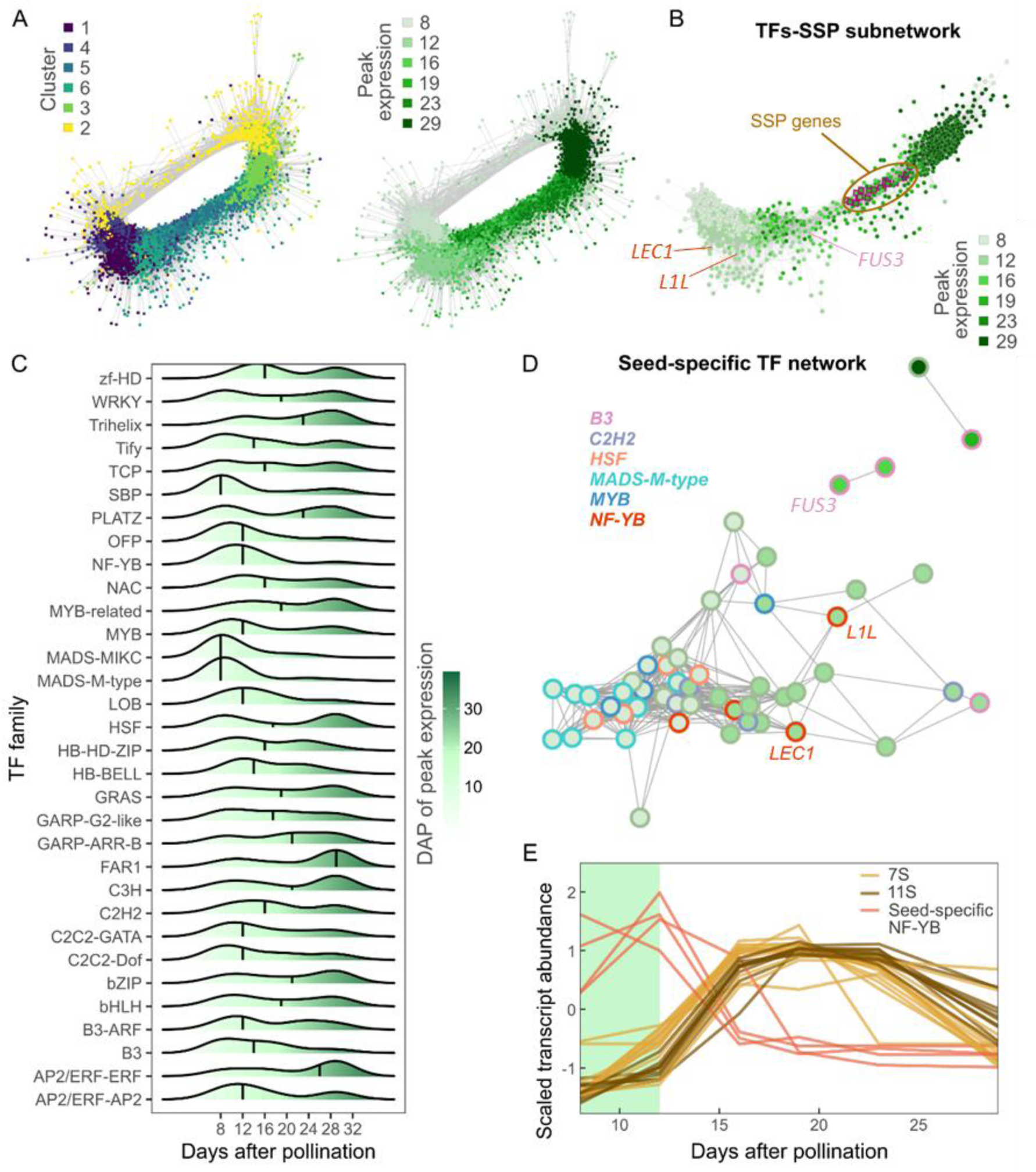
Temporal expression of transcription factors during pea seed development. (A) Co-expression network of differentially expressed genes across seed development. Connections between genes were identified using WGCNA and networks were visualized in Cytoscape using a Prefuse Force Directed layout, with an edge threshold cutoff of weight > 0.15. Nodes are colored by cluster (left) or by timing of peak expression (right). (B) Subnetwork of transcription factors (TFs) and seed storage protein (SSP, corresponding to 7S- and 11S-globulins) genes, with nodes colored by timing of peak expression. (C) Ridgeline plots representing the distribution of timing of peak expressions by TF family. Families with at least five members were considered. Vertical black lines indicate medians. (D) Subnetwork of seed-specific TFs. Families with at least three members are highlighted. (E) Transcript profiles of 7S- and 11S-globulins and of seed-specific NF-YB genes. Data are means of three biological replicates and are scaled. The green area indicates the embryogenesis seed developmental phase that ends at the 12 DAP stage (*i.e.* switch towards seed filling).

### *PsLEC1-like* is the major embryo-specific LEC1-type transcription factor in pea seed

Given the central role of the seed-specific NF-YB genes of the LEC1 type in controlling seed development and maturation in Arabidopsis (Jo et al., 2019), we examined the evolutionary relationships among the 19 NF-YB genes annotated in the V2 pea genome. Phylogenetic reconstruction based on NF-YB protein sequences from Pea, *A. thaliana* and *M. truncatula* revealed that two pea genes, *PsLEC1* (Psat.cameor.v2.3g82700) and *PsLEC1-like (PsL1L)* (Psat.cameor.v2.4g332150), clustered within the LEC1-type clade (Figure 3A). These two genes were in the seed-specific TF network in Figure 1D. Multiple sequence alignments further showed that both PsLEC1 and PsL1L harbored the conserved aspartate residue (corresponding to amino acid 55 in AtLEC1), a hallmark of LEC1-type NF-YB proteins required for seed-specific functions in *Arabidopsis* (Cagliari et al., 2014) (Figure 3B and Supplementary Figure S2A). At the whole-protein level, AtLEC1 and PsLEC1 shared 60% amino acid identity, whereas AtL1L and PsL1L shared 63%. This conservation increased markedly within the B domain, reaching 95.6 % identity between PsL1L and AtL1L (Supplementary Figure S2B). Expression profiling during seed development revealed that *PsL1L* is strongly expressed, with a peak at the transition between embryogenesis and seed filling (12 DAP), whereas *PsLEC1* follows a similar temporal pattern but at substantially lower levels (Figure 3C). Tissue-specific expression analyses by RT-qPCR across the three compartments of developing seeds (embryo, endosperm, and seed coat) showed that both genes were predominantly expressed in the embryo, with no detectable expression in the seed coat and only minimal levels in the endosperm (Figure 3D). Notably, *PsL1L* expression remained strongly expressed in the embryo at early stages of seed filling (Figure 3D), whereas *PsLEC1* displayed a similar localization pattern, but with a markedly lower level of expression (about 13-fold lower at 12 DAP). Altogether, these results support that *PsLEC1* and *PsL1L* are the respective orthologs of *AtLEC1* and *AtL1L*, and identify *PsL1L* as the major seed-specific LEC1-type gene in pea, consistent with its substantially higher expression relative to *PsLEC1*.

**Figure 3.**
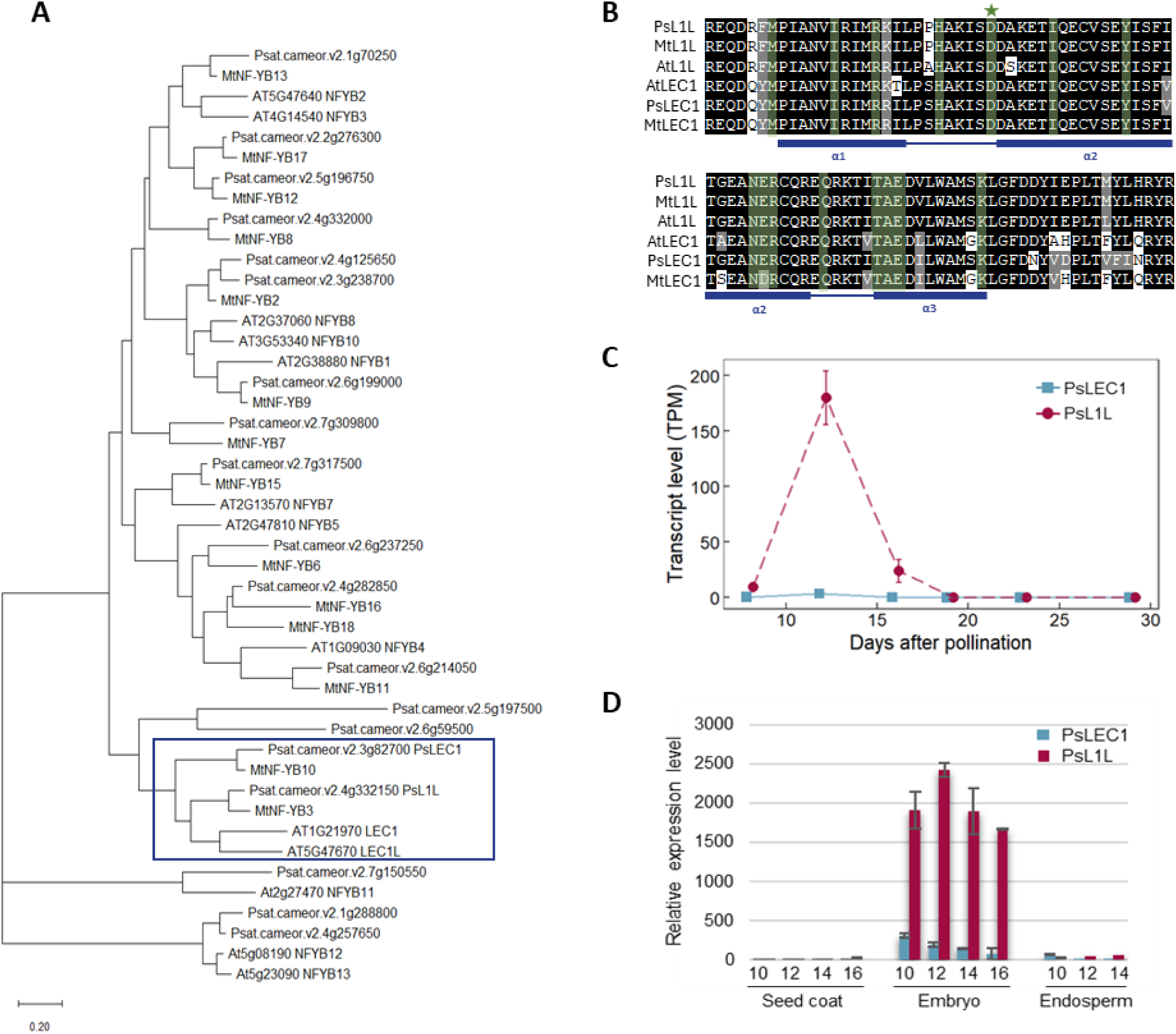
Phylogenetic analysis, B-domain conservation and expression patterns of LEC1-type NF-YB factors in pea. (A) Phylogenetic analysis of NF-YB transcription factors in pea (*Pisum sativum*), *Medicago truncatula*, and *Arabidopsis thaliana*. The LEC1/L1L clade is highlighted in blue. (B) Alignment of the conserved B domain, characteristic of the NF-YB family. Residues highlighted in black and gray represent identical and similar amino acids, respectively. Conserved residues shared between LEC1 and LEC1-ike but not in other Arabidopsis NF-YB proteins are highlighted in green. The aspartic acid residue, essential for AtLEC1 activity (Lee et al., 2003), is marked with a green star. Secondary structures (solid blue rectangles for alpha-helices and solid blue lines for coils) are represented based on Laloum et al., 2013. (C) Transcript (RNA-Seq) profiles of *PsLEC1* and *PsL1L* during seed development. Data are presented as means ± SD of four biological replicates. (D) Tissue-specific qRT-PCR analysis of *PsLEC1* and *PsL1L* expression in dissected seed tissues (embryo, endosperm and seed coat) collected from 10 to 16 days after pollination (DAP). At 16 DAP, the endosperm could not be collected as it had degenerated. Transcript levels were normalized to the reference genes *PsEF1α, PsACTIN* and *PsHKG1*. Data are presented as means ± SD of three biological replicates.

### Identification and molecular characterization of pea TILLING mutants for *PsLEC1-like*

To investigate *PsL1L* function during pea seed development, an EMS-mutagenized TILLING population (cv Caméor; Dalmais et al., 2008) was screened for mutations in *PsL1L*. A nonsense mutation introducing a premature stop codon at position 67, designated *l1l-1*, was identified and predicted to generate a truncated protein lacking the conserved B domain, thus likely non-functional (Figure 4A and Supplementary Figure S2A). Additional screening identified several missense mutations, including a non-synonymous amino-acid substitution at the beginning of the B domain changing a Valine residue into a Methionine (V85M) (Figure 4A and Supplementary Figure S2A), designated *l1l-2.* The impact of the mutations on the synthesis or stability of the *PsL1L* transcripts was studied in 12 DAP seeds of both alleles and their corresponding wild-type siblings, corresponding to the peak of expression of *PsL1L*. RT-qPCR analysis revealed a strong reduction in *PsL1L* transcript accumulation in *l1l-1* mutant seeds (Supplementary Figure S3), strongly suggesting that the premature stop codon caused nonsense-mediated decay of the mRNA (Popp and Maquat, 2016) as already observed for several pea TILLING non sense mutations (Vernoud et al., 2021; Bachelet et al., 2024). By contrast transcript abundance remained unchanged for the mRNA *l1l-2* variants (Supplementary Figure S3). Next, in order to evaluate the transcriptional activity of PsLEC1-like ad the functionality of the TILLING variants, we used a transient expression system in moss protoplasts established by Thévenin et al., (2016). In this system TF are expressed alone or in combination together with an *Arabidopsis OLEOSIN1* promoter fused to GFP (*pOLE1::GFP*), and GFP accumulation therefore serves as a quantitative proxy for promoter transactivation activity. Previous work demonstrated that LEC2 activates *pOLE1::GFP*, whereas LEC1 alone has no effect on *OLE1* promoter activity but enhances LEC2-mediated activation when co-expressed (Baud *et al*., 2016). Figure 5 shows the results obtained following co-expression of AtLEC1, PsL1L, or the TILLING variants, either alone or in combination with AtLEC2. As expected, AtLEC1 enhanced LEC2-mediated transcriptional activation of *pOLE1::GFP* but was inactive when expressed alone. PsL1L wild-type protein and the *L1L-2* missense variant displayed a similar behavior, both enhancing AtLEC2-mediated activation to an extent comparable to AtLEC1 (Figure 5). In contrast, co-expression of the *L1L-1* nonsense variant with AtLEC2 resulted in activation levels indistinguishable from those observed with AtLEC2 alone, indicating that the truncated protein is unable to potentiate LEC2-dependent transcriptional activation.

**Figure 4.**
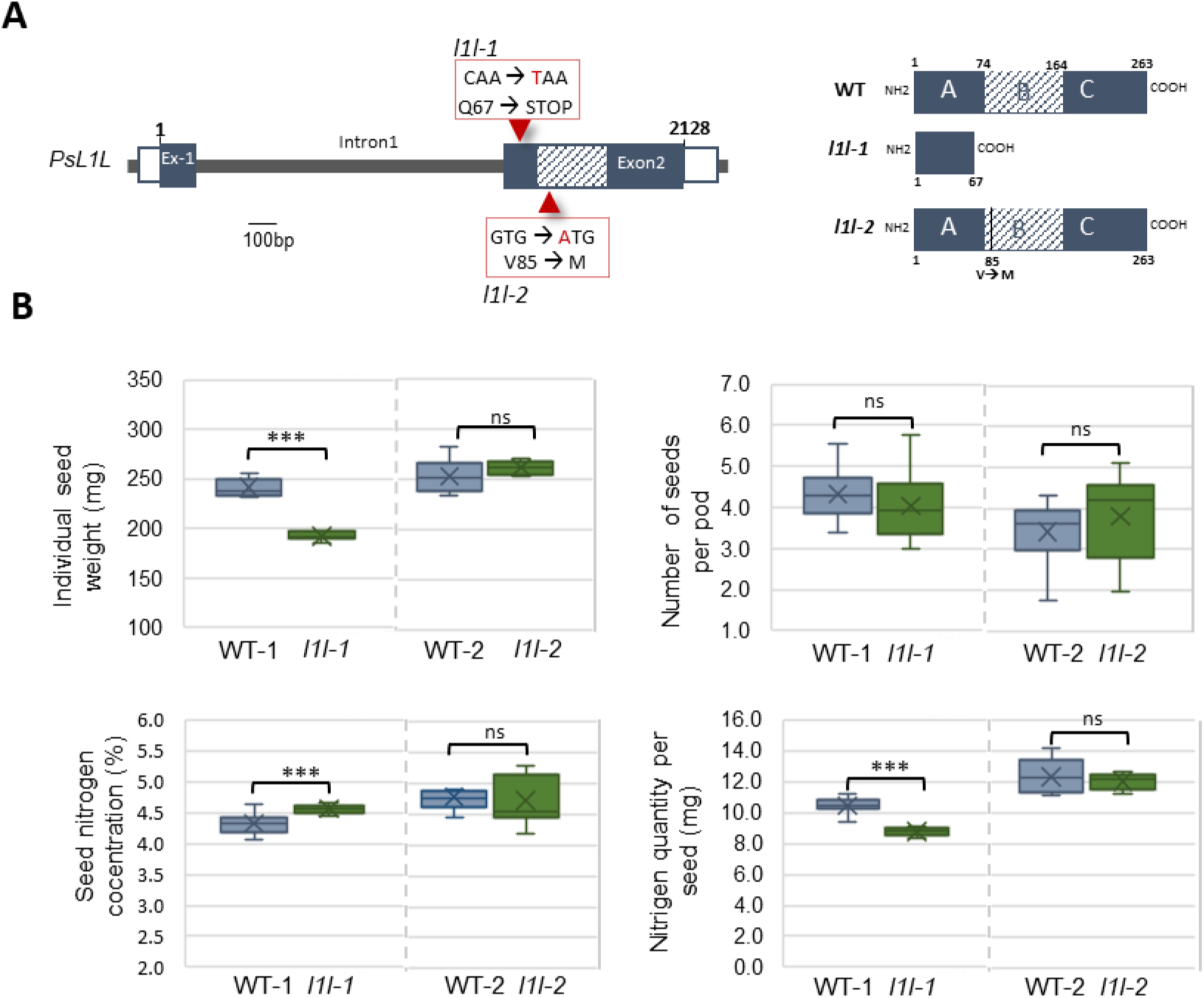
Loss of *PsL1L* function affects seed size and nitrogen accumulation. (A) Schematic representation of the *PsL1L* gene showing the positions of the TILLING mutations. The Q67stop mutation (*l1l-1* allele) introduces a premature stop codon before the conserved B domain, leading to a truncated protein (right panel) containing only the first 67 amino acids of the A domain. The V85M mutation (*l1l-2* allele) corresponds to a missense substitution at the beginning of the B domain. (B) Individual seed weight, number of seeds per pod, seed nitrogen concentration and absolute nitrogen quantity per seed (in mg) of the *l1l* TILLING alleles harvested at maturity compared to their respective wild-type (WT). Data represent means ± SD (n = 9 plants per genotype). Statistical significance was assessed using student’s t-test *** p<0.001, ns: not significant.

**Figure 5.**
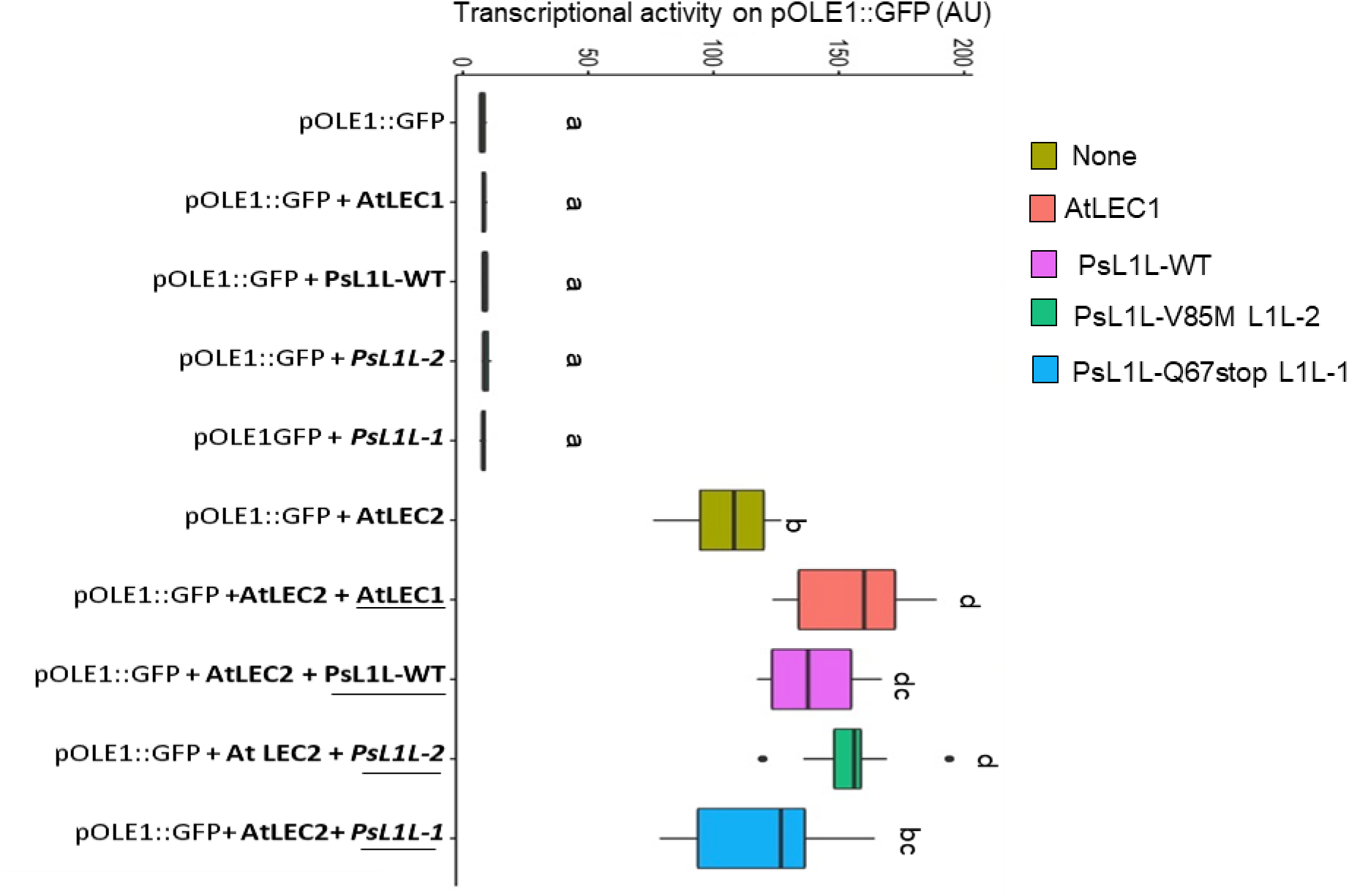
Transcriptional activity of PsL1L WT and TILLING-derived variants in moss protoplasts, with or without AtLEC2. GFP activity was quantified in moss (*Physcomitralla pattens*) protoplasts transfected with the *ProOLE1:*:*GFP* reporter together with constructs encoding AtLEC2, AtLEC1, PsL1L or two TILLING-derived variants: *PsL1L-1* introducing a premature stop codon at position 67 (Q67*), and *PsL1L-2*, carrying an amino acid substitution in the B domain (V85M). Constructs were expressed either individually or in combination with AtLEC2. Mean fluorescence intensity (AU, arbitrary units) was measured by flow cytometry. The *pOLE1::GFP* vector alone was used as a fluorescence control. Means of 9 replicates ± SD are presented. Groups sharing different letters differ significantly (Kruskal–Wallis test, α = 0.05; Dunn’s post hoc test with Benjamini–Hochberg correction).

### Loss of function of PsLEC1-like reduces seed size and affects early embryo growth

We next evaluated the effect of the mutations on mature seed traits by comparing each TILLING mutant with its corresponding segregating wild-type siblings (named WT1 and WT2 for *l1l-1* and *l1l-2* respectively, Figure 4B). The nonsense *l1l-1* allele caused a marked reduction in individual seed weight (−20%), while the number of seeds per pod was not significantly affected. This reduction in seed size was accompanied by a decrease in the absolute quantity of N per seed (−16%), even though seed N concentration showed a slight but significant increase (Figure 4B). By contrast, the missense *l1l-2* allele showed no significant differences compared with wild-type seeds, for any of the seed traits analysed (Figure 4B). In order to elucidate the developmental basis of the reduced seed size observed in the loss-of-function mutant, we further characterized the *l1l-1* allele through a seed growth kinetics analysis. WT and mutant seeds were collected from 8 to 38 DAP (every 4–5 days), spanning embryogenesis, the transition phase, seed filling, and late maturation (Figure 6A). A significant reduction of one-seed fresh and dry weights was observed in the *l1l-1* mutant compared to the wild type from the transition phase onward (12 DAP) and persisting throughout seed filling until late maturity (38 DAP, seed water content <50%) (Figure 6A). In contrast, seed water content was not significantly affected (Figure 6A), indicating that seed developmental timing was not altered, and that the phenotype more likely reflects impaired seed growth rather than a delay in seed development. An independent second experiment covering embryogenesis (6–10 DAP) and early seed filling (up to 20 DAP; Supplementary Figure S4) confirmed these results and further showed that the size differences appeared at 10 DAP, consistent with the high expression of *Ps1L1* at this stage. Overall, these results indicate that *PsL1L* is required for normal seed growth from the late embryogenesis onward, without affecting developmental timing.

**Figure 6.**
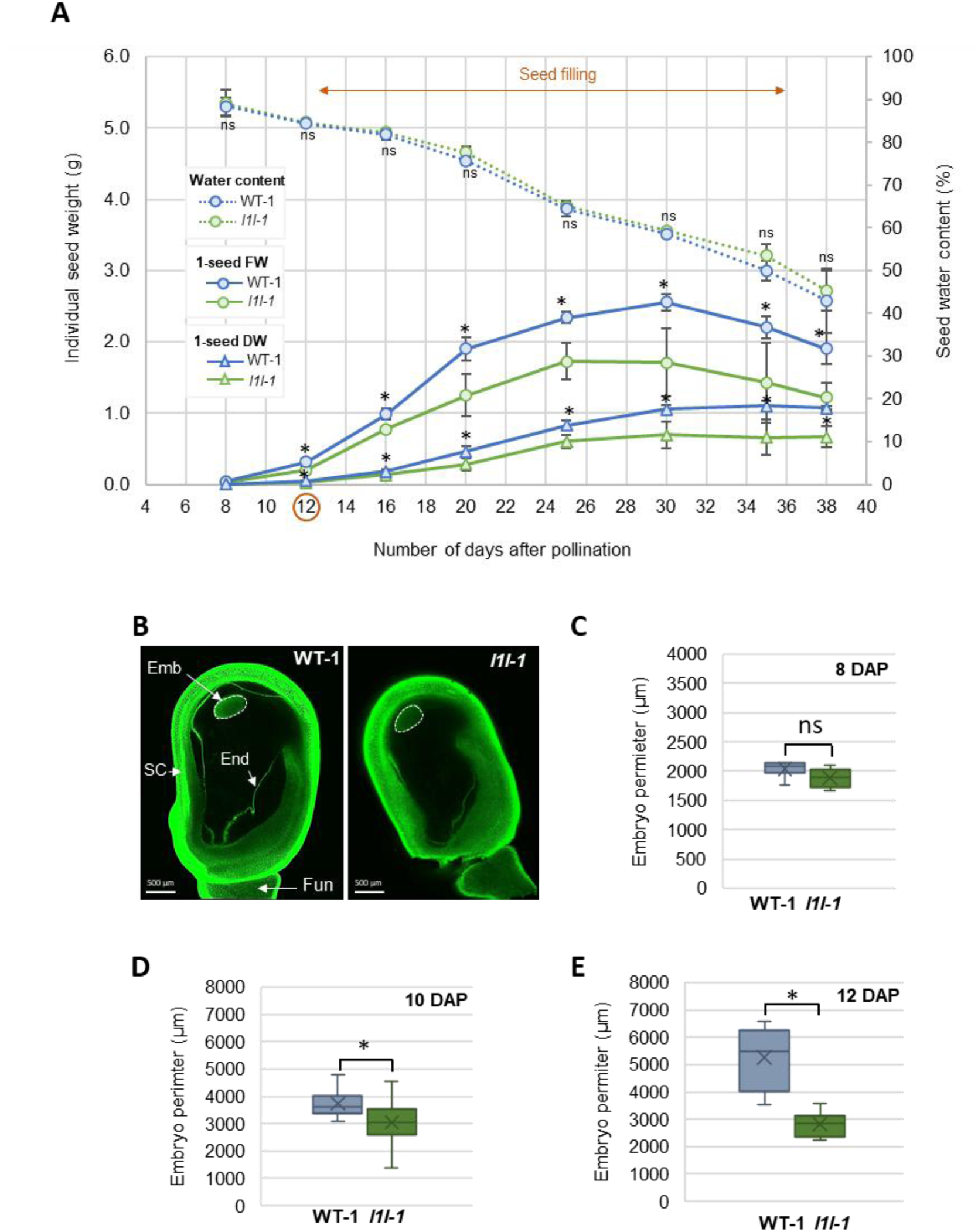
*PsL1L* controls embryo and seed growth. (A) Kinetics of seed development in wild-type (WT) and *l1l-1* loss of function allele. Individual seed fresh weight, dry weight and seed water content were measured from 8 to 38 days after pollination. Differences in seed weight between WT and mutant seeds appear early at the transition phase (12 DAP), and persist throughout seed filling. Data represent means ± SD (n = 6-8 pods per genotype and time point, see material and methods). (B) Representative confocal images of BABB-cleared wild-type (WT-1) and *l1l-1* seeds at 8 DAP. Autofluorescence generated by glutaraldehyde fixation allows visualization and measurement of the embryo proper within intact seeds. Emb, embryo; End, endosperm; SC, seed coat, Fun, funiculus (C-E) Embryo perimeter measured from cleared seeds at 8, 10 and 12 DAP. Data represent means ± SD of 6 individual seeds. Statistical significance was assessed using the Mann–Whitney test; *P<0.05; ns : not significant.

During early seed development, maternal tissues play a predominant role in determining seed size, as the embryo remains relatively small, prior to its rapid growth phase during seed filling. This prompted us to assess whether the reduced seed size observed in the mutant during embryogenesis was also associated with defects in embryo development. Embryogenesis was examined in the *l1l-1* loss-of-function mutant using an adapted benzyl alcohol/benzyl benzoate (BABB) tissue clearing protocol optimized for pea seeds and enabling high-resolution confocal imaging and embryo measurements (Figure 6B, see Materials and Methods). At 8 DAP, embryo size did not differ significantly between mutant and wild type (Figure 6C), consistent we the seed weight measurements (Figure 7A). However, *l1l-1* embryos were significantly smaller at 10 and 12 DAP (Figure 6, D and E). Altogether these results reveal *PsL1L*-dependent control of early embryo and seed expansion.

**Figure 7.**
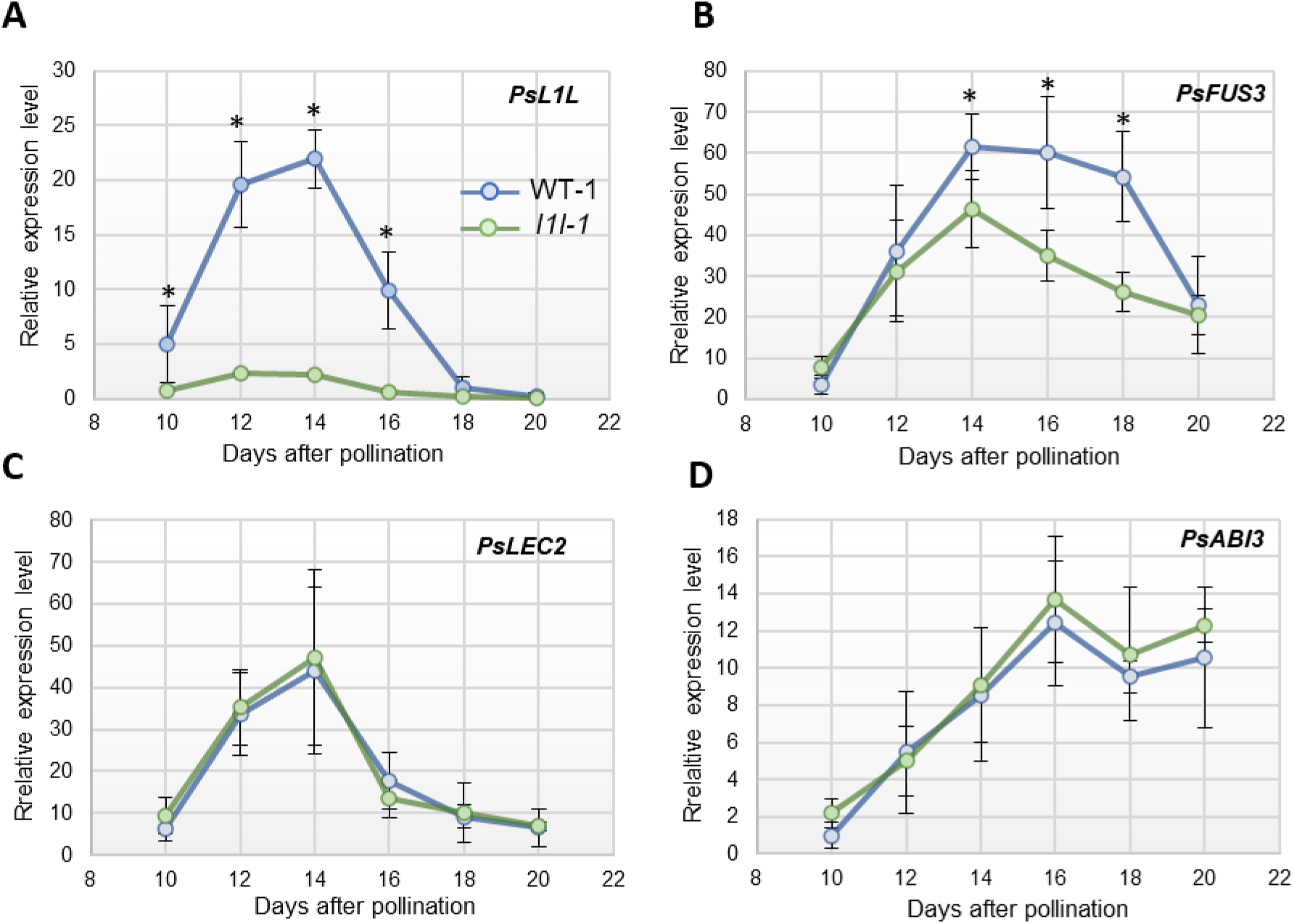
Expression profiles of the AFL genes during seed development in *WT and l1l-1* mutant seeds. qRT-PCR analysis of (A) *PsL1L, (B) PsFUS3 (Psat.cameor.v2.3g165200)*, (C) *PsLEC2 (Psat.cameor.v2.7g221600)*, (D) *PsABI3 (Psat.cameor.v2.3g90900)* in WT and *l1l-1* developing seed from 10 to 20 days after pollination. Transcript levels were normalized to the reference genes *PsEF1α* and *PsHKG1*. Data represent means ± SD of four biological replicates. Statistical significance was assessed using the Mann–Whitney test; * P<0.05.

### *PsL1L* influences the expression of *PsFUS3* but not that of *PsLEC2* ad *PsABI3*

In *Arabidopsis*, LEC1 orchestrates multiple developmental processes, including embryogenesis, by regulating distinct sets of genes, notably the AFL B3-domain TFs LEC2, ABI3, and FUS3 (Jo et al., 2019). To investigate whether a similar regulatory relationship exists in pea and whether PsL1L affects the LAFL seed regulatory network, we quantified the expression of these TFs during seed development in the *l1l-1* mutant and WT seeds from 10 to 20 DAP (Figure 7), covering the period of *PsLEC1-like* expression and its subsequent decline (Figure 7A). Phenotypic data from this developmental time-course experiment are presented in Supplementary Figure S4. *PsL1L* expression was strongly reduced in the mutant seeds throughout the analysed period (Figure 7A), confirming the results obtained at 12 DAP (Figure S3). In WT seeds, *PsLEC2* (Psat.cameor.v2.7g221600) displayed an expression pattern similar to that of PsL1L, whereas *PsFUS3* (Psat.cameor.v2.3g165200) showed a later expression peak between 14 and 18 DAP. *PsABI3* (Psat.cameor.v2.3g90900) exhibited a distinct profile, with expression beginning at 16 DAP and persisting throughout seed filling. Interestingly, *PsFUS3* expression was significantly downregulated in *l1l-1* mutant seeds between 14 and 18 DAP, whereas the expression kinetics of *PsABI3* and *PsLEC2* were not affected (Figure 7 B-D).

## Discussion

In this study, we generated a comprehensive seed expression atlas covering six developmental stages in pea (*Pisum sativum* cv Cameor), providing a detailed temporal resolution of transcriptional dynamics throughout seed development in this species. Using this resource, we identified several seed-specific TFs, including PsL1L belonging to the NF-YB gene family, which is highly expressed during the key transition from embryogenesis to seed filling, and further highlighted its role in seed size control.

Our transcriptomic analysis, conducted using the recently released improved pea genome assembly (Kreplak et al., 2026), revealed extensive transcriptional reprogramming accompanying pea seed development, with nearly 80% of expressed genes showing significant temporal variation across the developmental stages examined. Clustering analysis revealed transcriptional modules corresponding the canonical developmental phases previously described in legumes: embryogenesis, seed filling and late maturation (Figure 1). The latter was related to cluster 3 with genes peaking at 29 DAP, it was enriched in genes involved in protein turnover, RNA processing and responses to water deprivation ((Figure 1 and Supplemental Dataset3), consistent with the acquisition of desiccation tolerance and metabolic quiescence, as well as mRNA accumulation for protein synthesis during germination. On the other hand, genes from cluster 1 and 4 were preferentially expressed during early stages of seed development and were enriched in functions related to DNA replication, chromatin organization, cytoskeleton dynamics and cell wall biogenesis, reflecting intense cell proliferation and embryo growth (Figure 1 and Supplemental Dataset3). Similar functional enrichments have been reported in developing seeds of other legume species (Thompson et al., 2009; Yao et al., 2016; Kudapa et al., 2018).

Cluster 4 was mainly composed of genes showing peak expression at 8 DAP, and in our seed-specific TFs network composed of 52 TF (Figure 2D, Dataset S2and S4), 14 belonged to this cluster. Several members of the MADS-box M-type family were represented (Figure 2C), including homologs of *AtAGL62* (*AGAMOUS-like 62*) (Psat.cameor.v2.6g299550 and Psat.cameor.v2.6g299650; Supplementary Dataset S2). In *Arabidopsis thaliana*, *AGL62* is specifically expressed in the endosperm and has been shown to prevent premature cellularization during the syncytial phase of endosperm development (Kang et al., 2008). Loss of *AtAGL62* function leads to seed lethality associated with extremely early endosperm cellularization. In pea, the endosperm initially develops as a coenocyte, similarly to *Arabidopsis* (Marinos, 1970). However, the extent of endosperm cellularization remains unclear. The pea endosperm is generally described as “liquid” up until the point at which it disappears, and the cellularization process itself is still poorly documented and understood (Marinos, 1970, Melkus et al., 2009). The functional characterization of key regulators such as *AGL62* homologs could provide important insights into these processes.

Clusters 6 and 5 defined two successive transcriptional phases during seed filling, corresponding respectively to early (16-19 DAP) and late (19-23 DAP) seed filling stages (Figure 1). Genes from cluster 6 were enriched in functions related to photosynthesis, carbohydrate homeostasis, nitrogen responses and lipid metabolism (Figure 1 and Supplementary Dataset3), highlighting the strong coordination between metabolic activity and reserve synthesis during this phase. Developing legume embryos are known to undergo a photoheterotrophic metabolism in which embryonic photosynthesis contributes to oxygen production and energy supply, thereby supporting biosynthetic fluxes and reserve accumulation during seed filling (Weber et al., 2005; Rolletschek et al., 2003). Genes from cluster 5 contained several major SSP genes, including 7S and 11S globulins annotated as cupin-like proteins, as well as lectins, lipoxygenases and LEA (late abundant embryogenesis) proteins, all known to accumulate to high levels in mature legume seeds. The enrichment of the GO term “response to virus” in this cluster (Figure 1C) reflects the presence of these cupin and lectin family proteins, which are frequently associated with defense-related annotations/GO terms (Supplementary Dataset S3). Cluster 5 also contained 83 TFs (Supplementary Dataset S2), several of which were directly connected to SSP genes in our TF-SSP subnetwork (Figure 2C; Supplementary Dataset S5). Among them, *Psat.cameor.v2.5g67950*, the pea homolog of *AtbZIP67*, was directly connected to several 11S globulin genes (Supplementary Dataset S4). In Arabidopsis, AtbZIP67 is involved in the regulation of seed storage compound accumulation together with other bZIP TF and members of the LAFL complex (Bensmihen et al., 2003; Yamamoto et al., 2009; Kim et al., 2022). Consistent with this, several B3-domain TFs, including FUS3 (Psat.cameor.v2.3g165200), were also identified in the TF-SSP network, in line with the presence of RY cis-elements in pea globulin promoters (Kreplak et al., 2019) mediating the seed-specific expression of legumin genes (Bäumlein et al., 1992).

Analysis of our transcriptomic data identified also major reprogramming during the switch form embryogenesis to seed filling, occurring when seed water content is below 85% (Ney et al., 1998) that is at 12 DAP in our experimental set up. Understanding the processes underlying the embryogenesis/filling transition is of particular interest as it might help to modulate both seed size and storage capacities. In legumes, this switch is characterized by metabolic and developmental changes, notably the differentiation of the embryo cotyledon epidermis into transfer cells necessary for nutrient uptake (Weber et al., 2005). Members of the HD-ZIP IV transcription factor that control differentiation of epidermal cell types in plants were identified in our TF network, (Schrick et al., 2023). In pea, differentiation of cotyledon transfer cells is a key determinant of the transition from embryogenesis to seed filling, as shown by mutant studies (Borisjuk et al. 2002), but the underlying molecular mechanisms of this differentiation remain poorly understood. Whether HD-ZIP IV TFs contribute to this process will be an interesting avenue for future investigation. Members of the NF-YB family were also identified in the seed-specific network (Figure 2 C-E). Given the important role of seed-specific NF-YB genes in regulating seed maturation and development, we further investigated the function and evolutionary relationships of these genes.

Based on phylogenetic reconstruction, sequence identity, and expression profiling, we identified *PsLEC1* (Psat.cameor.v2.3g82700) and *PsLEC1-like (PsL1L)* (Psat.cameor.v2.4g332150) as the respective orthologs of AtLEC1 and AtLEC1-like. Both pea proteins possess the conserved amino acid residues characteristic of the LEC1 clade, including the aspartic residues located at position 55 in Arabidopsis, which is critical for AtLEC1 function during embryogenesis (Lee et al., 2003). *PsLEC1* and *PsL1L* are expressed early during seed development, in both the embryo and the endosperm, as shown in Arabidopsis (Jo et al., 2019). However, *PsL1L* exhibits substantially higher expression levels than *PsLEC1* from the end of embryogenesis onward, with a peak at the transition stage that persists into early seed filling (Figures 2E, 3CD and 7A). Consistent with this expression pattern, the PsLEC1-like protein, but not PsLEC1, was quantified by shotgun proteomics during the transition stage and early seed filling in pea (Henriet et al., 2021), further supporting a predominant role for this seed-specific NF-YB TF during this developmental phase.

In order to better understand the role of *L1L* in pea we screened the Caméor TILLING population for mutant in this gene. The *l1l-1* allele carries a premature stop codon producing a truncated protein devoid of both the highly conserved B domain and the C domain. Transient expression assays in moss confirmed that the full length *PsL1L* protein is functional and is able to increase transcriptional activity of LEC2 on a seed specific promoter, in a similar way as AtLEC1 (Figure 5). The loss of activity observed for the truncated L1L-1 protein in this system, together with the dramatic decrease in its mRNA abundance *in planta* (Figure 7A and S3), confirm that this allele is a true loss-of-function mutation, which explains the developmental and metabolic phenotypes observed in *l1l-1* seeds. In contrast, the *l1l-2* V85M variant retained transcriptional activity in moss in the presence of AtLEC2 (Figure 5), and showed no modification in its mRNA abundance (Figure S3) consistent with its lack of seed phenotype (Figure 4B). These results reinforce the idea that *PsL1L* activity depends on the formation of the canonical NF-Y complex, as has been show with Arabidopsis *LEC1* and *L1L* (Calvenzani et al., 2012). They also indicate that the absence of domains B and C in PsL1L, as for AtLEC1 (Lee et al., 2003) severely impaired NF-YB function in seeds

Functional analysis of the *l1l-1* allele revealed that L1L plays a crucial role in determining seed size in pea. Mature mutant seeds were significantly smaller and contained lower nitrogen amount than WT seeds (Figure 4). Kinetic analyses of seed development, together with imaging of cleared embryos, demonstrated that *l1l-1* seeds and embryos were already smaller at 10-12 DAP. These observations suggest a defect in early embryo expansion rather than a simple delay in the seed developmental program, consistent with the absence of differences in seed water content, a recognized marker of seed developmental progression in legumes (Ney et al., 1993). Early embryo growth is a major determinant of final seed size in legumes, as it establishes the cellular capacity that subsequently limits the accumulation of storage reserves (Gallardo et al., 2007). Reduced embryo growth in the mutant is therefore likely to limit the capacity of cotyledons to accumulate storage compounds, including proteins, consistent with the decrease in seed N content observed in *l1l-1* seeds (Figure 4B). In *Arabidopsis*, the role of LEC1 has been extensively characterized, with mutants displaying multiple developmental defects affecting embryogenesis and reserve accumulation (reviewed in Jo et al., 2019). In contrast, the function of *AtL1L* remains less well understood. Initial studies using an RNAi construct driven by the 35S promoter showed that suppression of *AtL1L* expression induced embryo developmental defects distinct from those observed in *lec1* mutants (Kwong et al., 2003). Subsequently, the characterization of two *AtL1L* T-DNA insertion lines did not reveal altered phenotypes during seed development, and no effect on mature seed size was reported (Yamamoto et al., 2009). It was suggested that the absence of phenotype in *Arabidopsis l1l* mutants may result from functional redundancy with AtLEC1, which remains fully expressed in these lines (Yamamoto et al., 2009). Our results suggest that this is not the case in pea and provide, to our knowledge, the first evidence that an LEC1-LIKE factor plays a direct role in controlling embryo growth and final seed size in legumes. These observations position as a regulator that influences seed yield potential by controlling embryo growth capacity, more similar to the one described for AtLEC1. However, although *PsL1L* is able to complement AtLEC1 in our transactivation assay in moss (Figure 5), the absence of a strong seed phenotype similar to At*lec1* in the l1l-1 TILLING line suggests nevertheless a different role in the process than At*LEC1*.

In *Arabidospsis*, LEC1 acts upstream of ABI3, FUS3, and LEC2, as ABI3 and FUS3 expression is reduced in *lec1* mutants, while LEC1 overexpression leads to increased ABI3 and FUS3 expression in developing seeds (Kagaya et al., 2005; To et al., 2006; Pelletier et al., 2017). Moreover, *AtABI3*, *AtFUS3*, and *AtLEC2* are directly transcriptionally regulated by AtLEC1 (Pelletier et al., 2017). Interestingly, our data show that loss of PsL1L specifically reduces *PsFUS3* transcript levels, while *PsLEC2* and *PsABI3* remain largely unaffected. *FUS3* is known to regulate both embryo development and storage protein gene expression, and therefore represents a likely downstream mediator of PsL1L function as suggested in Arabidopsis (Kagaya et al., 2005; Wang & Perry, 2013). Consistent with this, our shotgun proteomics data of early seed development (Henriet et al., 2021) indicates a temporal succession between these factors, with PsLEC1-like detected earlier than PsFUS3 during seed development, supporting the hypothesis that PsL1L functions upstream of PsFUS3 to promote embryo growth and storage reserve initiation.

## Supplemental data

**Figure S1.**
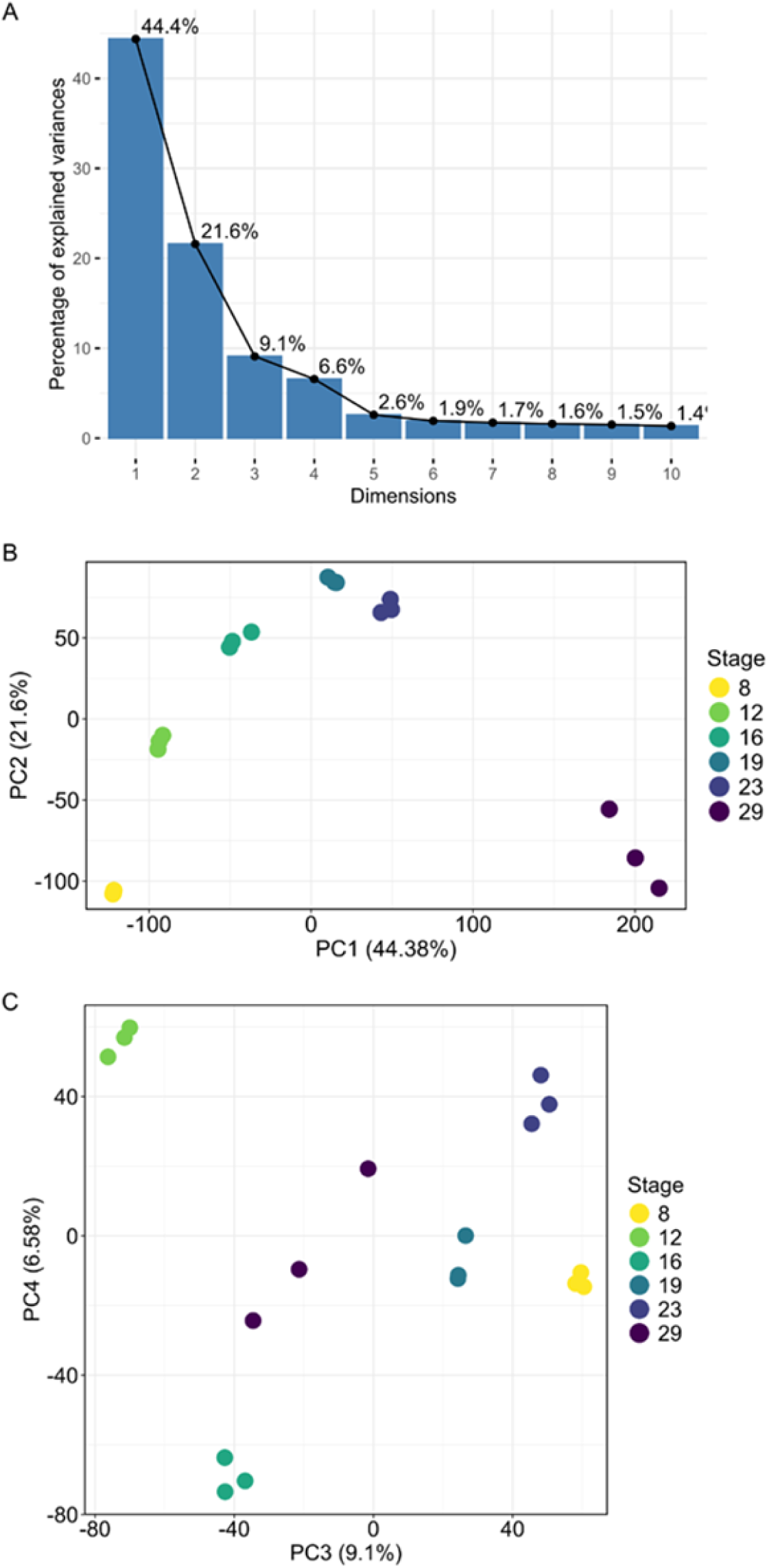
Principal Component Analysis (PCA) of the developing pea seed transcriptome. (A) Barplot showing the percentage of explained variance of the first 10 principal components. (B) and (C), PCA plots highlighting separation of samples on PC1 and PC2 (B) and on PC3 and PC4 (C).

**Figure S2.**
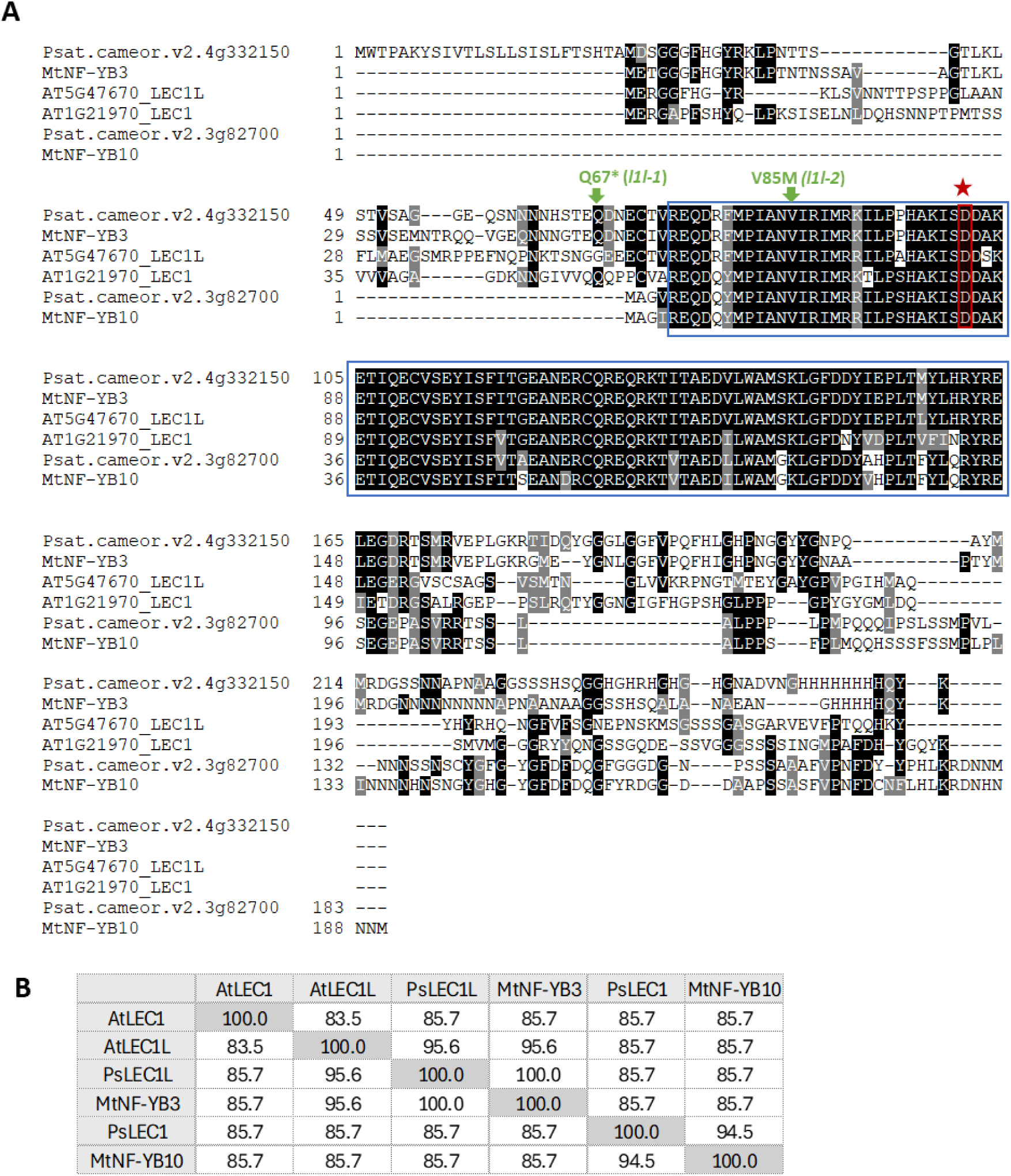
Sequence alignment and conservation of LEC1 and LEC1-like proteins. (A) Multiple protein sequence alignment of LEC1 and LEC1-like proteins from pea (*Pisum sativum*), *Arabidopsis thaliana*, and *Medicago truncatula*. The highly conserved B domain, characteristic of the NF-YB subunit family, is highlighted in blue. The aspartic acid residue, essential for LEC1 activity (Lee et al., 2003), is highlighted in red and marked with a red star. Green arrows indicate the EMS-induced point mutations identified by TILLING. (B) Percentage identity matrix of the conserved B domain among LEC1 and LEC1-like proteins. Values represent pairwise amino acid sequence identity (%).

**Figure S3.**
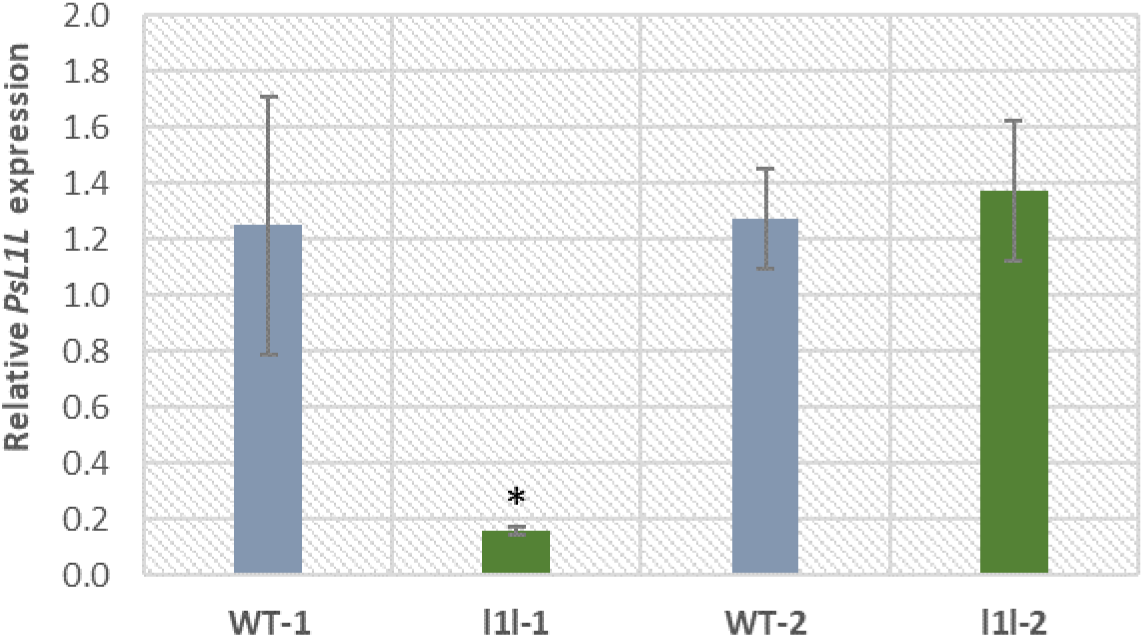
Relative *PsL1L* expression in 12 DAP seeds of TILLING mutants and their respective wild-type. *PsL1L* expression was quantified by RT-qPCR and normalized to the reference genes *PsEF1α* and *PsACTIN*. Values were further normalized to WT-1. *PsL1L* expression was reduced by approximately 8-fold in the *l1l-1* mutant compared with its wild-type control. Values represent the mean ± SD of three biological replicates. Asterisks indicate statistical significance (Mann-Whitney test, p < 0.05).

**Figure S4.**
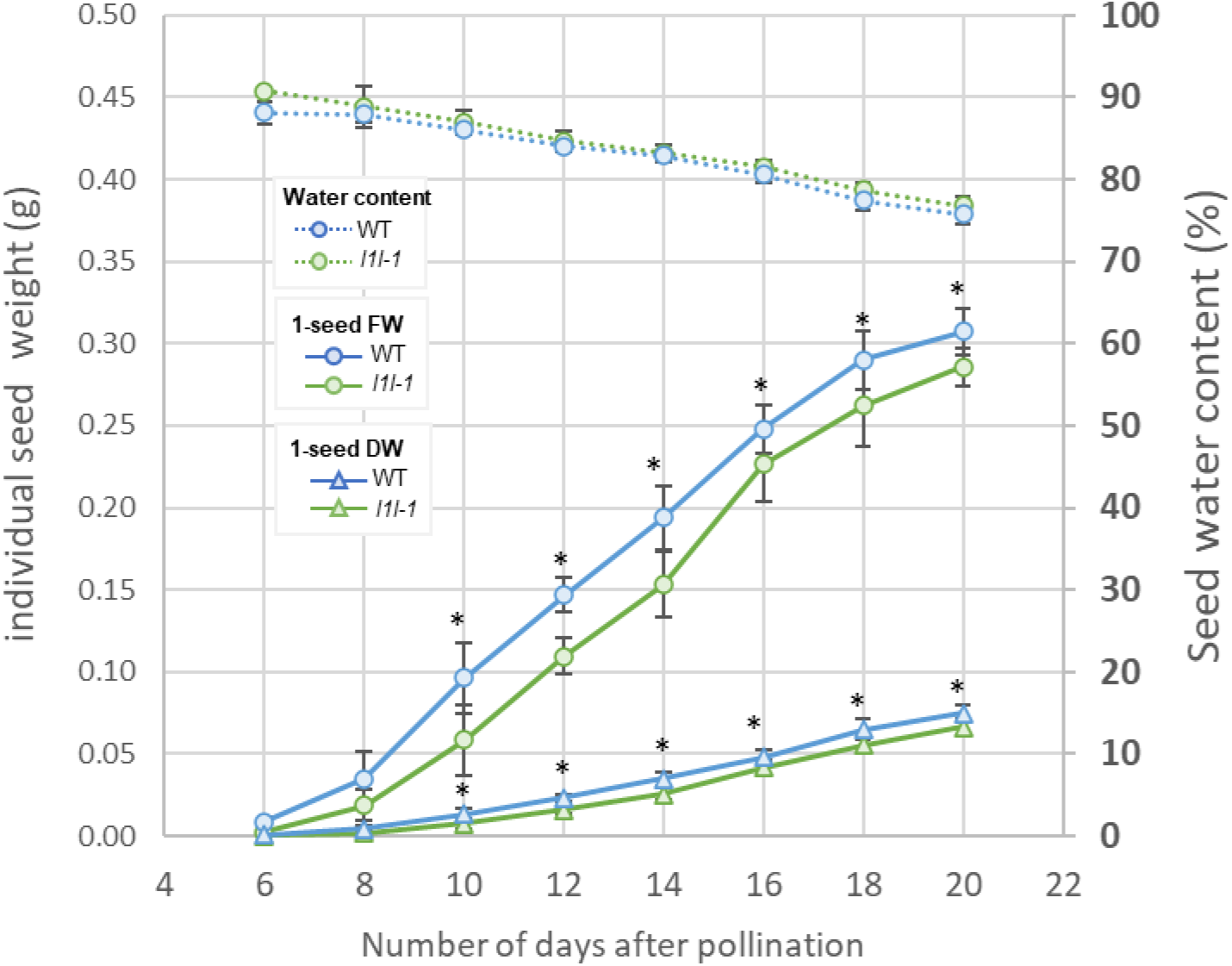
Kinetics of seed development in wild-type and *l1l-1* mutant from 6 to 20 days after pollination. Developing wild-type (WT) and mutant pods were harvested every two days from 6 to 20 days after pollination. For each genotype and developmental stage, 4 to 15 pods were collected, opened on ice, and seeds were retrieved, counted, and used for fresh weight (FW), dry weight (DW; 80 °C for 2 days), and calculation of mean individual seed FW, DW, and water content using the formula [(FW − DW) / FW] × 100. Data are presented as means ± SD. Statistical significance was assessed using the Mann–Whitney test; *p<0.05. Seeds from the same experiment were also collected from 10 to 20 DAP for RNA extraction and RT-qPCR analyses presented in Figure 7.

**Table S1.**
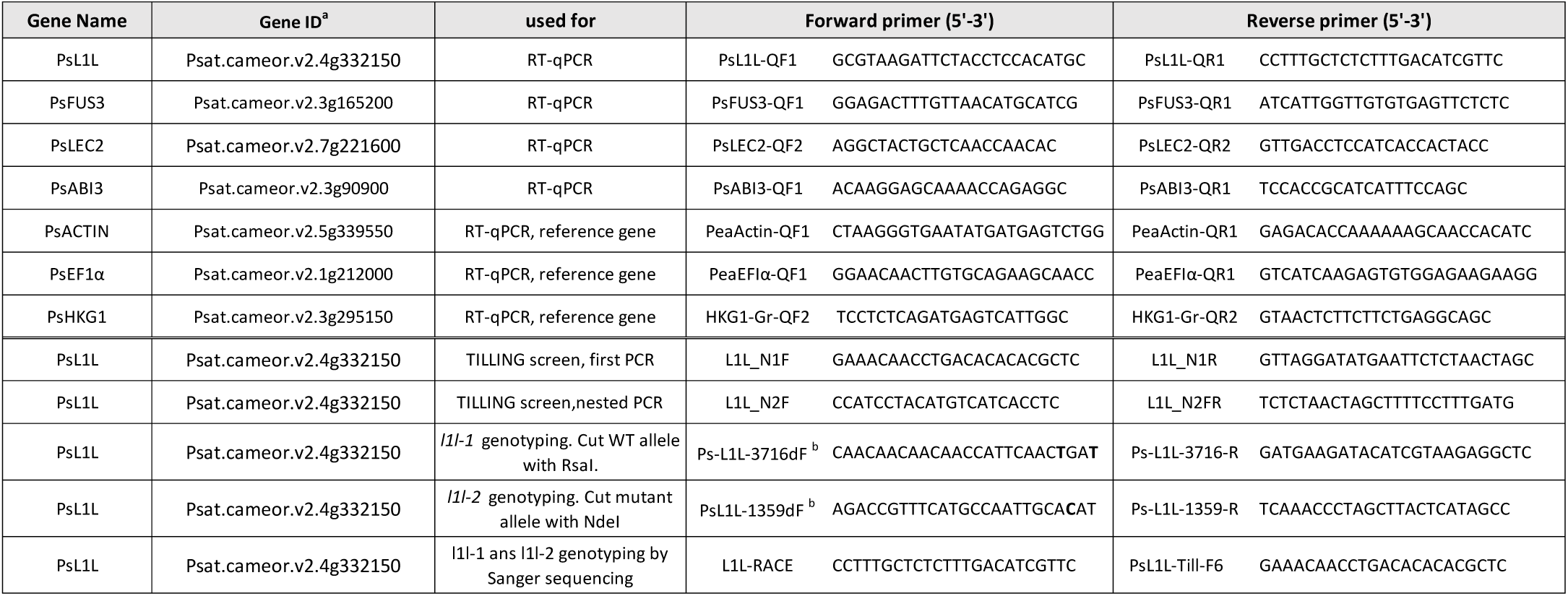
Primers used in this study. ^a^ Gene ID are from the V2 Caméor version (Kreplak et al., 2026). ^b^ The CACGACGTTGTAAAACGAC tail was added to the 5’ end of the dCAPS primer; the nucleotide corresponding to the introduced mutation is shown in bold.

**Table S2.** Phenotypic data of seeds used for RNA-seq transcriptomic analysis. Values are mean ± SD of n=4 biological repeats. DAP, days after pollination. FW: fresh weight; DW: dry weight.

| DAP | 1-seed FW (g) | 1- seed DW (g) | Water content (%) | Seed developmental stages |
| --- | --- | --- | --- | --- |
| 8 | 0.02 ± 0.004 | 0.003 ± 0.001 | 86.9 ± 0.51 | Embryogenesis |
| 12 | 0.128 ± 0.027 | 0.021 ± 0.005 | 83.7 ± 0.67 | Transition |
| 16 | 0.28 ± 0.022 | 0.058 ± 0.006 | 79.1 ± 0.49 | Filling |
| 19 | 0.389 ± 0.0445 | 0.117 ± 0.015 | 69.9 ± 0.93 | Filling |
| 23 | 0.466 ± 0.043 | 0.175 ± 0.018 | 62.4 ± 1.29 | Filling |
| 29 | 0.4446 ± 0.048 | 0.218 ± 0.017 | 50.7 ± 2.35 | Maturation |

**Supplementary Dataset S1:** Normalized gene expression in developing pea seeds.

**Supplementary Dataset 2.** Clustering of differentially expressed genes in developing pea seeds.

**Supplementary Dataset 3.** Enriched GO terms in clusters of differentially expressed genes in developing pea seeds.

**Supplementary Dataset 4.** Pea transcription factors: identification of seed-specific expressed TF.

**Supplementary Dataset 5.** Edge weight of connected nodes in the TF-SSP gene network.

## Acknowledgments and Fundings

We thank the members of the 4PMI platform (Phenotyping Platform for Plant–Microorganism Interactions, INRAE, Dijon, France) for their technical support for plant growth. We acknowledge Rémy Félix-Serre and the GeT-PlaGe core facility (Toulouse, France) for RNA-seq library preparation and sequencing. We thank Laure Avoscan and Chrystel Deulvot from the DImaCell microscopy platform (INRAE/University of Burgundy, Dijon, France) for helpful discussions, advice, and availability. We also thank Katia Belcram from Institut Jean-Pierre Bourgin for Plant Sciences (Versailles, France) for her advice on BAAB clearing. We are grateful to Sirine Benmamar for her work on the moss transactivation assay and to Clélia Picard for providing the qPCR results shown in Figure S3 and her help for the kinetic experiment from Figure 6. This work was supported by the French National Research Agency (ANR) through the REGULEG project (ANR-15-CE20-0001) and the PEAVALUE project (ANR-19-CE21-0008).

